# LRRK2 kinase inhibition protects against Parkinson’s disease-associated environmental toxicants

**DOI:** 10.1101/2024.03.29.587369

**Authors:** Neda M. Ilieva, Eric K. Hoffman, Mohammed A. Ghalib, J. Timothy Greenamyre, Briana R. De Miranda

**Affiliations:** Center for Neurodegeneration and Experimental Therapeutics, Department of Neurology, University of Alabama at Birmingham, Birmingham, AL, USA; Pittsburgh Institute for Neurodegenerative Diseases, Department of Neurology, University of Pittsburgh, Pittsburgh PA, USA

**Keywords:** Parkinson’s Disease (PD), Gene x Environment (GxE), Environmental Toxicants, Leucine Rich Repeat Kinase 2 (LRRK2), Mitochondria

## Abstract

Idiopathic Parkinson’s disease (PD) is epidemiologically linked with exposure to toxicants such as pesticides and solvents, which comprise a wide array of chemicals that pollute our environment. While most are structurally distinct, a common cellular target for their toxicity is mitochondrial dysfunction, a key pathological trigger involved in the selective vulnerability of dopaminergic neurons. We and others have shown that environmental mitochondrial toxicants such as the pesticides rotenone and paraquat, and the organic solvent trichloroethylene (TCE) appear to be influenced by the protein LRRK2, a genetic risk factor for PD. As LRRK2 mediates vesicular trafficking and influences endolysosomal function, we postulated that LRRK2 kinase activity may inhibit the autophagic removal of toxicant damaged mitochondria, resulting in elevated oxidative stress. Conversely, we suspected that inhibition of LRRK2, which has been shown to be protective against dopaminergic neurodegeneration caused by mitochondrial toxicants, would reduce the intracellular production of reactive oxygen species (ROS) and prevent mitochondrial toxicity from inducing cell death. To do this, we tested *in vitro* if genetic or pharmacologic inhibition of LRRK2 (MLi2) protected against ROS caused by four toxicants associated with PD risk – rotenone, paraquat, TCE, and tetrachloroethylene (PERC). In parallel, we assessed if LRRK2 inhibition with MLi2 could protect against TCE-induced toxicity *in vivo*, in a follow up study from our observation that TCE elevated LRRK2 kinase activity in the nigrostriatal tract of rats prior to dopaminergic neurodegeneration. We found that LRRK2 inhibition blocked toxicant-induced ROS and promoted mitophagy *in vitro*, and protected against dopaminergic neurodegeneration, neuroinflammation, and mitochondrial damage caused by TCE *in vivo*. We also found that cells with the LRRK2 G2019S mutation displayed exacerbated levels of toxicant induced ROS, but this was ameliorated by LRRK2 inhibition with MLi2. Collectively, these data support a role for LRRK2 in toxicant-induced mitochondrial dysfunction linked to PD risk through oxidative stress and the autophagic removal of damaged mitochondria.

## 1. Introduction

Environmental contaminants such as the pesticides rotenone^1–5^ and paraquat ^1,4,6,7^, as well as the commonly used organic solvent trichloroethylene (TCE)^8–12^ increase Parkinson’s disease (PD) risk by approximately 0.7 – 2.5-fold^4,8,9^. Given the similar risk for neurodegeneration from structurally distinct environmental toxicants, convergent mechanisms on cellular pathways such as mitochondrial damage and subsequent oxidative stress are implicated in the pathogenesis of idiopathic PD^13–19^. We and others have previously shown that systemic exposure to environmental mitochondrial toxicants, such as rotenone, paraquat, and TCE, cause the selective degeneration of nigrostriatal dopaminergic neurons^20–22^. In addition, experimental models of exposure to these toxicants result in oxidative stress, neuroinflammation, deficits in the autophagy lysosomal pathway (ALP), and the accumulation of alpha-synuclein (αSyn) within dopaminergic neurons or cells in the surrounding ventral midbrain, all of which are pathologies commonly found in postmortem PD brain tissue^23–25^.

In addition to toxicant-induced brain pathology, growing experimental evidence indicates that a common mechanism of neurotoxicity caused by environmental exposures may involve the protein **<u>L</u>**eucine-**<u>R</u>**ich **<u>R</u>**epeat **<u>K</u>**inase 2 (LRRK2), an autosomal dominant inherited risk factor for PD^22,25,26^. LRRK2 is a multi-domain kinase protein with both GTP-ase and kinase activity, which act to phosphorylate Rab GTP-ases^27^. These GTP-ases function in the maturation of vesicles in the endolysosomal and vesicular trafficking pathways and regulate many cellular homeostatic mechanisms such as the packaging of cell materials and autophagy and mitophagy^28,29^. The G2019S mutation, the most commonly inherited LRRK2 mutation associated with PD risk, is a toxic gain-of-function mutation located in the kinase domain, which causes oxidative damage, impairments in mitophagy and autophagy, as well as vesicular trafficking dysfunction^30,31^. Data have shown that inhibition of LRRK2 kinase activity, such as with the pharmacological inhibitors PF360 (Pfizer) or MLi2 (Merck), can decrease aberrantly activated LRRK2 associated deficits in cellular functions such as membrane trafficking and autophagy and mitophagy^18,32^. While the function and pathology of elevated mutant LRRK2 kinase activity has been shown consistently in G2019S mutant models, G2019S mutations in human populations are incompletely penetrant for PD, with approximately 28% elevated risk for disease at age 59, and 79% elevated risk at age 79^33,34^. This incomplete penetrance underscores the important contributing risk of environmental factors, which could influence PD development and progression.

Aberrant activation of wildtype (WT) LRRK2 is also suggested to play a role in idiopathic PD, as elevated LRRK2 kinase activity has been observed in brain tissue and urinary exosomes and PBMCs of individuals with PD with no known genetic mutations^25,26^. We and others have shown that environmental toxicants associated with PD risk, such as rotenone and TCE, elevate WT LRRK2 kinase activity in rodent brain tissue^1,35^. We also found that TCE exposure elevated WT LRRK2 kinase activity in the brain of rats to similar levels as G2019S LRRK2 knock-in mice, and this elevated kinase activity preceded nigrostriatal dopaminergic neuronal degeneration and coincided with oxidative damage within surviving dopaminergic neurons^23^. As LRRK2 is a major regulator of autophagy and mitophagy function^32,35–41^, we suspected that elevated LRRK2 kinase activity may play a role in toxicant-induced oxidative stress by impairing mitophagy or turnover of damaged mitochondria. As dopaminergic neurons are highly vulnerable to mitochondrial dysfunction^42^, the convergence of environmental toxicants and LRRK2 at the mitochondria could explain the selective degeneration of dopaminergic neurons from toxicant exposure. Conversely, this may demonstrate a mechanism for how LRRK2 kinase inhibition is protective against mitochondrial toxicants.

To assess this, we developed an *in vitro* model using WT, LRRK2-G2019S, or LRRK2 KO human embryonic kidney (HEK) cells to measure the role of LRRK2 in reactive oxygen species (ROS) and mitophagy following exposure to rotenone, paraquat, TCE, and a structurally similar chlorinated solvent, tetrachloroethylene (PERC). We complemented our *in vitro* findings using MLi2 to block LRRK2 kinase activity in a rat model of TCE exposure that we previously demonstrated elevated nigrostriatal LRRK2 kinase activity^22^. We found that LRRK2 inhibition significantly protected against toxicant-induced oxidative stress and mitochondrial damage, both in cultured cells and in nigrostriatal dopaminergic neurons, possibly as a result of restored mitophagy, which was ultimately neuroprotective. Using LRRK2 knockout cells, we further confirmed that inhibition of LRRK2 was sufficient to protect against environmental toxicant-induced oxidative stress, which was exacerbated in the G2019S mutation. Collectively, these data show a potential role for LRRK2 kinase activity in neurodegeneration from mitochondrial toxicants, and how therapeutic treatment with LRRK2 kinase inhibitors could be protective against neurotoxicity induced by environmental exposures associated with PD.

## 2. Materials and Methods

### 2.1. Animals, TCE Exposure, and MLi2 Treatment

All procedures were performed in compliance with relevant laws and institutional guidelines and have been approved by the Institutional Animal Care and Use Committee at the University of Alabama at Birmingham (UAB). Twelve-month-old female Lewis rats were obtained through a retired breeding program through Envigo (Indianapolis, IN.) At arrival, rats were separated into double housing and allowed to acclimate for two weeks before beginning treatment. Thirty rats were randomly distributed into 3-week and 6-week groups, with ten and twenty rats in each group respectively. Conventional diet and water were maintained ad libitum, with animals remaining in a temperature-controlled environment with a 12-hour light-dark cycle. Female rats were chosen due to their mass and the availability of MLi2 dosed at 10 mg/kg over 3 weeks via oral gavage. We previously published no sex differences have been observed at the level of neurodegeneration or LRRK2 kinase activity in animals exposed to TCE^22,43^.

Trichloroethylene (TCE, ThermoFisher) was dissolved into premium olive oil (Trader Joe’s, Monrovia, CA) for a final concentration of 200 mg/kg based on individual rodent weight. MLi2 (synthesized by MedChemExpress) was dissolved into a 1% methyl cellulose solution to a final concentration of 10 mg/kg based on individual body weight. Three-week rats were randomly assigned as either vehicle (VEH; olive oil) or TCE treatment (n = 5 rats each). Six-week rats were randomly assigned to either vehicle or TCE treatment with or without MLi2 (n = 5 rats each). Animal weights were recorded weekly to adjust the dosing of TCE and MLi2 to maintain appropriate concentrations. Animals were euthanized at either the 3-week or 6-week endpoints respectively with a lethal dose of pentobarbital, followed by transcardial perfusion with 1x phosphate buffered saline (PBS) with phosphatase and protease inhibitors. Fresh brains were sectioned in half, with half allotted for post-perfusion fixation, and half were frozen immediately at -80°C. Post-perfusion fixation was performed with 4% paraformaldehyde for 24 hours, and then brains were switched to 30% sucrose until saturated for sectioning on a freezing microtome.

### 2.2 CRISPR/Cas9 gene edited HEK 293-T cells (G2019S, LRRK2 KO)

CRISPR/Cas9 LRRK2 mutants were generated using the approach described by Di Maio et al^26^. Briefly, to generate the LRRK2 KO cell line, CRISPR/cas9 editing was modified to targeted exon41 of LRRK2. A specific guide RNA (5′-ATTGCAAAGATTGCTGACTAGTTTT-3′) was cloned into a GeneArt CRISPR Nuclease Vector. HEK-293T cells were transfected with this vector via nucleoporation using a Nucleoporater II device, followed by collection and FACS enrichment. Sorted cells were then subsequently interrogated using exon-41 targeted PCR amplification and DNA sequencing to confirm the genotype. G2019S knock-in cell lines were generated similarly but consisted of transfection of 120-mer single-stranded oligonucleotide containing a G to A substitution, which codes for the glycine-to-serine amino acid change.

### 2.3 Immunohistochemistry

Brains sectioned with a microtome at 35 μm with a 1/6 serial fraction. Tissue was preserved in cryoprotectant at -20°C immediately after sectioning for future immunostaining. For immunofluorescent immunohistochemistry, brain sections were washed in 1x PBS, then blocked with 10% Normal Donkey Serum (NDS; Jackson Immunoresearch) in 1x PBS-0.3% Triton for one hour. Tissue was blocked in primary antibodies in 1% NDS in 1x PBS-0.3% Triton (**Table 1**) overnight at 4°C on a shaker, then washed in 1x PBS, and blocked in AlexaFluor secondary antibodies for one hour under dark conditions. The tissue was washed, and then DAPI (1:5000) was applied for 5 minutes. The tissue was then washed again before being wet-mounted and coverslipped. All fluorescent microscopy images were obtained using a Nikon AX Confocal System with 60x oil Nikon Lambda series objective. All images were analyzed using Nikon Elements AR 5.9 software.

For 3,3’-Diaminobenzidine (DAB) staining, tissue was washed in 1x Tris Buffered Saline (TBS) before quenching with 0.6% H_2_O_2_ in methanol for 20 minutes. Sections were then subjected to heat-based antigen retrieval with citrate buffer (10 mM sodium citrate with 0.05% Tween, pH 6.0) for 30 min at 30°C. After, sections were washed, before blocking with 10% Normal Goat Serum (NGS; Jackson ImmunoResearch) in 1x TBS-0.3 % Triton for one hour at room temperature on a shaker. Tissue was then incubated in anti-Rabbit Tyrosine Hydroxylase (TH) (1:2000; EMD Millipore) in 1% NGS in 1x TBS-0.3% Triton overnight at 4°C with agitation. Tissue was then washed, before a 4-hour incubation in 1:500 biotinylated Goat anti-Rabbit IgG (H + L) (Vector Labs) secondary in 2.5% NGS in 1x TBS-0.3% Triton at room temperature, with agitation. The tissue was then washed again before incubation in ABC solution (Vector Labs) in 1x TBS for 30 minutes on a shaker. DAB substrate chromogen solution (Vector Labs) was then used for staining, followed by washing and wet mount. Tissue was dried on the slides, before dehydration. Sections were then coverslipped with Permount and left to dry overnight before imaging. Images were imaged using an MBF brightfield microscope, and stereology counts were completed using StereoInvestigator 5.1 Software for objective volumetric stereology counts. Brightfield images were obtained at 4x air objective on an MBF brightfield microscope.

### 2.4. HEK Cell Oxidative Stress Measurement

Wildtype 293-T HEK cell lines were CRISPR-edited to express the LRRK2 G0219S mutation or with complete LRRK2 knockout at the University of Pittsburgh as previously described in Keeney et al., 2021^44^. Cell lines were cultured in complete Dulbecco’s Modified Eagle Medium (standard DMEM, Fetal Bovine Serum (FBS), and Penicillin-Streptomycin; ThermoFisher). Cells were plated at 0.05 x 10^6^ seeding density into Poly D Lysine-coated glass-coverslipped 24-well plates and grown until about 70% confluency then treated for 4 or 24 hours with Dimethylsulfoxide (DMSO; 500 μM) with or without MLi2 (1 μM), Rotenone (ROT; 500 nm), Paraquat (PQ; 100 μM), trichloroethylene (TCE; 500 μM), or tetrachloroethylene (PERC; 500 μM), in complete DMEM (500 μL per well). Cells were then either incubated with Dihydroethidium (DHE) dye (3 mM DHE in DMSO at 1:1000) for 30 minutes prior to fixation or washed with PBS and fixed immediately with 4% Paraformaldehyde in PBS with Magnesium-Chloride (250 μL per well) for 30 minutes. DHE-dyed cells were then counterstained with DAPI (1:5000) in PBS for 5 minutes, and then washed and mounted on standard microscope slides. DHE-dyed cells were imaged the following day on a Nikon AX microscope with a 60x oil Nikon Lambda series objective. DHE signal was quantified by measuring the mean intensity of the TRITC channel after drawing bezier ROIs around cells, N = 3-5 independent experimental biological replicates.

### 2.4. Immunocytochemistry

After fixation, HEK 293-T cells and respective mutant clones (LRRK2-G2019S and LRRK2 KO) were washed three times with 1x PBS before blocking with 10% Normal Donkey Serum in PBS + 0.02% Triton for 1 hour at room temperature with agitation. Cells were then incubated with primary antibodies (**Table 1**) in 1% NDS in PBS + 0.3% Triton overnight at 4°C. Cells were then washed three times the next day, followed by secondary incubation with AlexaFluor secondaries in 1x PBS at room temperature with agitation for 1 hour under dark conditions. Coverslips were washed again, before staining with DAPI (1:5000) in PBS for 5 minutes and washed twice more. Coverslips were then mounted inversely onto standard microscope slides with mounting media and kept in the dark until imaging. Imaging was performed using a Nikon AX Imaging system using 60x oil lambda-series objectives. Analyses were performed using Nikon Elements AR 5.4 software. DHE mean intensity was quantified via ROIs per image (N = 3 – 5 independent biological replicates). Number of intersecting LC3b/TOM20 puncta were analyzed per ROI using the binary intersection function in Nikon Elements software and were size restricted to only include puncta above 0.05 μM.

### 2.5. Statistical analyses

All data in bar graphs are expressed as mean values ± standard error of the mean (SEM). *A priori* power calculations were performed using G*power software (Heinrich-Heine-University Düsseldorf) based on prior data to determine the sample size required for a 20 – 40% difference between means for a 95% power with α set at 0.05. *P*-values are reported for the interaction comparison between “genotype” and “treatment”. All data was modeled and tested for normality using thorough normality testing for residuals, homoscedasticity, and variance before being analyzed using the appropriate statistical test. Statistical outliers from each data set were determined using the extreme studentized deviate (Grubbs’ test, α = 0.05) and were only analyzed if conditions determined *a priori* were met. *A priori* conditions for statistical exclusion evaluation included if a singular data point influenced each mean enough to affect significance, and if distribution was skewed and non-parametric. One-way and two-way ANOVA were performed where indicated, with post-hoc analyses for multiple comparisons performed with appropriate post-hoc tests dependent on number of comparisons. A standard t-test was used to compare 3-week treated vehicle and trichloroethylene *in vivo* animal groups. Statistical significance between treatment groups is represented in each figure as *p<0.05, **p<0.01, ***p< 0.001,***p<0.0001, unless otherwise indicated in figure legends. Statistical analysis was carried out using GraphPad Prism software (v10.2.0, GraphPad Software, Boston, MA).

### 2.6. Materials

**Table 1.** Antibody and reagent information for immunocytochemistry (ICC), immunohistochemistry (IHC) with either DAB (chromogenic), or immunofluorescence (IF) imaging.

| Assay | Antibody Name | Catalog Info | Company | Dilution |
| --- | --- | --- | --- | --- |
| ICC | Dihydroethidium (DHE) | D11347 | Thermo Fisher | 1:1000 |
|  | DAPI | 28718-90-3 |  | 1:5000 |
|  | Rabbit LC3b Polyclonal | PA5-35195 | Invitrogen | 1:1000 |
|  | Mouse TOM20 [4F3] | ab56783 | Abcam | 1:1000 |
| IHC (DAB) | Rabbit TH | AB152 | EMD Millipore | 1:2000 |
| IHC (IF) | Sheep TH | AB1542 |  | 1:2000 |
|  | NeuroTrace 640/660 Nissl | N21483 | Invitrogen | 1:500 |
|  | Rabbit 3-Nitrotyrosine | A21285 |  | 1:1000 |
|  | Mouse 4-Hydroxynonenal | MAB3249 | R&D Systems | 1:1000 |

All other reagents were purchased through Thermo Fisher Scientific (Waltham, MA) unless specifically stated.

## 3. Results

### 3.1. LRRK2 mediated PD-associated environmental toxicant-induced oxidative stress

To assess whether LRRK2 kinase activity has a functional role in mediating oxidative stress *in vitro*, HEK 293-T cells were selected as a well-characterized cell line with established quantities of LRRK2^45^. To evaluate the effect of LRRK2 genotype on toxicant-induced oxidative stress, WT 293-T, LRRK2-G2019S, and LRRK2 knockout (KO) cells were treated with either toxicants alone; rotenone (ROT), paraquat (PQ), TCE, or PERC, or in combination with 1 µM of the LRRK2 kinase inhibitor MLi2 (**Fig. 1A**). Quantitative analyses of DHE signal per cell demonstrated significant effects of the genotype, treatment, and interaction effects of genotype and treatment for each toxicant (ROT: p<0.0001; PQ: p<0.0001; TCE: p<0.0001; PERC: p<0.0001, two-way ANOVA; **Fig. 1B-E**). LRRK2 genotype significantly contributed to ROS expression, with G2019S cells showing elevated ROS at baseline compared to WT that was exacerbated by toxicant treatment (ROT: p<0.0001; PQ: p<0.0001, two-way ANOVA) while LRRK2 KO cells are significantly protected from ROS elevation across all toxicant treatments (ROT: p=0.0017; PQ: p<0.0001; TCE: p<0.0001; PERC: p<0.0001, two-way ANOVA). LRRK2 kinase inhibition with MLi2 significantly reduced oxidative stress in WT 293-T HEK cells and in LRRK2 G2019S gain of function mutants. Detailed ANOVA Summary Tables, descriptive statistics, and post-hoc multiple comparisons are provided in **Supplemental Table 1**.

**Figure 1.**
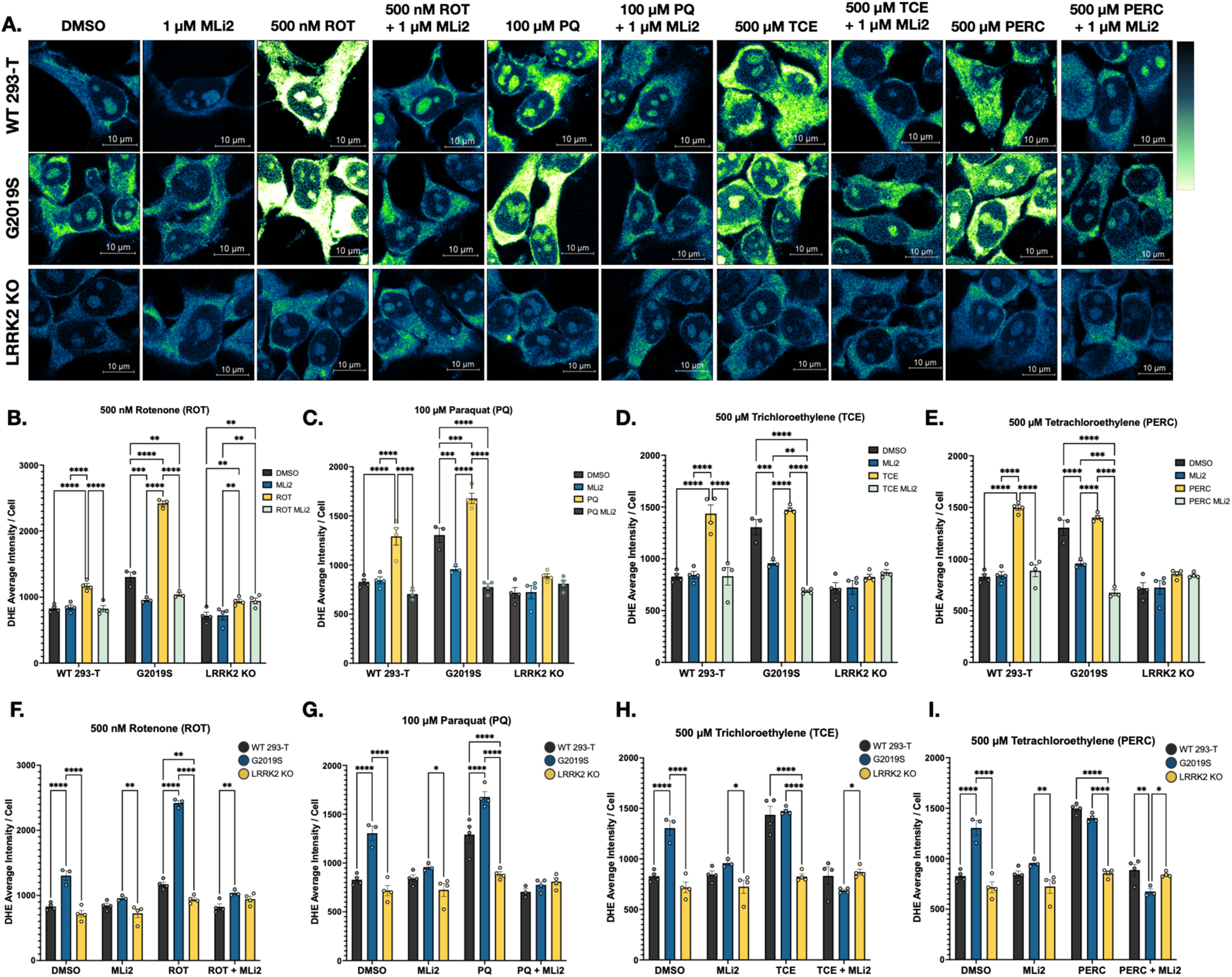
LRRK2 mediates toxicant induced ROS generation. **A** WT or CRISPR-edited HEK-293T cells (G2019S, LRRK2 KO) were treated with 500 nM ROT, 500 μM PQ, 1 mM TCE or PERC alone or in combination with 1 μM MLi2 as per previously identified time points (4 hours for ROT & PQ, and 24 hours for TCE & PERC). ROS were measured with dihydroethidium (DHE) and DHE signal is represented in a pseudocolored gradient (blue is lowest ROS and white is highest ROS).**B-E** DHE signal was increased in all toxicant-treated groups and exacerbated in G2019S cells. DHE was significantly decreased in the MLi2-treated groups and in LRRK2 KO cells. Significant treatment effects were found. ROT (F(3,32) = 135.4, p<0.0001); PQ (F(3,33) = 63.61, p<0.0001); TCE (F(3,34) = 47.79, p<0.0001); PERC (F(3, 33) = 74.59, p<0.0001). **F – I.** DHE signal was significantly elevated in G2019S cells compared to WT 293-T for ROT and PQ treatments, but not for TCE and PERC. MLi2 restored DHE signal to WT and LRRK2 KO levels in G2019S cells. Significant effects of genotype were found. ROT (F(2,32) = 188.0, p<0.0001); PQ (F(2,33) = 61.18, p<0.0001); TCE (F(2,34) = 39.12, p<0.0001); PERC (F(2, 33) = 56.93, p<0.0001). Significant interaction effects of treatment and genotype were found. ROT (F(6,32) = 50.55, p<0.0001); PQ (F(6,33) = 14.60, p<0.0001); TCE (F(6,34) = 18.59, p<0.0001); PERC (F(6, 33) = 26.65, p<0.0001). N = 3 – 5 biological replicates, 120 - 160 cells per technical replicate, two-way ANOVAs with post-hoc Tukey test for multiple comparisons adjustment. ANOVA tables, descriptive statistics, and post-hoc comparisons are provided in Supplemental Table 1. Bar graphs represent mean values for each group, with error bars. Significance values as indicated: **p<0.01, ***p<0.001, ****p<0.0001.

### 3.2. Toxicant-induced oxidative stress is mediated by LRRK2 in a threshold-dependent manner

To investigate the specificity of LRRK2 in oxidative stress caused by toxicant exposure, LRRK2 KO cells were treated with a dose-response of the prototypical mitochondrial complex I inhibitor ROT, and evaluated for ROS using DHE. While a 500 nM ROT dose did not produce a significant elevation in ROS in LRRK2 KO cells, doses at 1 μM ROT and beyond (2, 4, and 8 μM) caused significant ROS elevation compared to DMSO-treated cells (**Fig. 2A-B**; DMSO vs. 500 nM ROT: p = 0.2568; DMSO vs. 1, 2, 4, 8 μM ROT: p<0.0001; one-way ANOVA). This threshold between 500 nM and 1 μM ROT was then used in a follow-up experiment to evaluate the specificity of LRRK2 inhibition in attenuating oxidative stress (**Fig. 2C-D**). LRRK2 KO and WT 293-T cells were treated with 500 nM ROT or 1 μM ROT alone or in combination with 1 μM MLi2 and DHE signal was quantified and evaluated (**Fig. 2C-D, E-F** respectively). Both LRRK2 KO and WT 293-T cells displayed significantly elevated ROS with 500 nM ROT (LRRK2 KO p=0.0386; WT 293-T p=0.0054), however only WT 293T cells displayed reduced ROS with MLi2 treatment (LRRK2 KO p>0.9999, WT 293-T p=0.0048, one-way ANOVA respectively). Neither WT nor LRRK2 KO cells displayed reduced ROS with LRRK2 inhibition at 1 µM ROT (p=0.9868, and p=0.274, respectively, one-way ANOVA) indicating that oxidative stress caused by severe toxicity (near complete complex I inhibition) is likely non-LRRK2 mediated or overcomes the regulation of this system.

**Figure 2.**
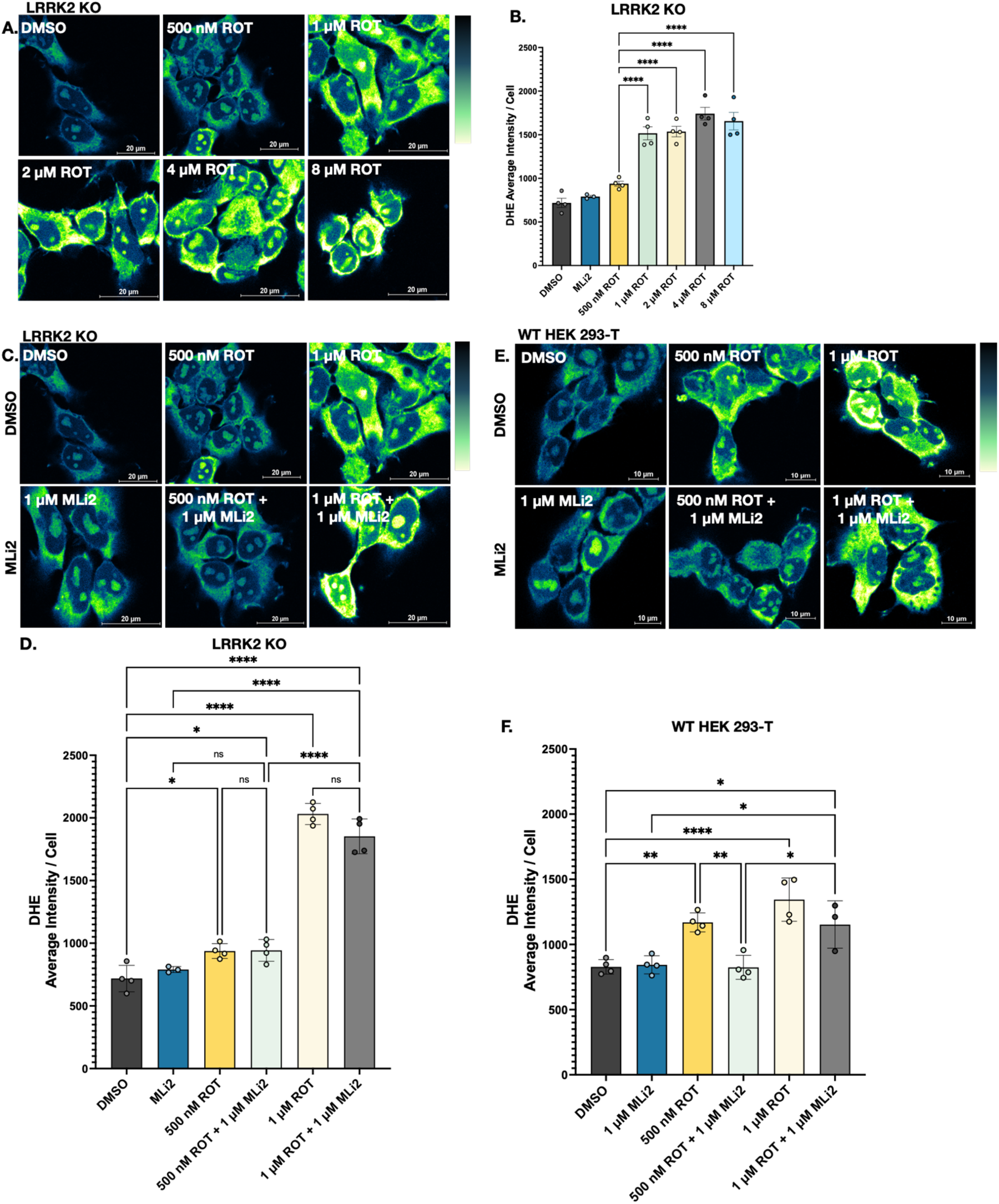
LRRK2 mediated effects on toxicant induced ROS are threshold dependent. **A.** Representative images of CRISPR-edited 293-T cells (LRRK2 knockout; LRRK2 KO) treated with either 0.10% DMSO, 1 μM MLi2, or scaled doses of rotenone (ROT; 500 nM, 1 μM, 2 μM, 4 μM, and 8 μM ROT). **B.** ROS quantification using DHE signal intensity of LRRK2 KO cells. ROS formation was significantly increased only in LRRK2 KO cells treated with more than 1 μM ROT (p<0.0001) with no significant differences between 1 – 8 μM ROT; (F(6,20) = 42.97, p<0.0001), one-way ANOVA. **C.** Representative images of LRRK2 KO cells treated with 0.10% DMSO, 1 μM MLi2, 500 nM ROT, or 1 μM ROT alone or in conjunction with 1 μM MLi2. **D.** ROS quantification using DHE signal intensity of LRRK2 KO cells. No significant differences were shown between 500 nM or 1 μM ROT alone or with MLi2 (p>0.9999, and p=0.1239 respectively). One-way ANOVA, (F(5,17) = 148.2, p<0.0001). **E.** Representative images WT 293-T cells treated with 0.10% DMSO, 1 μM MLi2, 500 nM ROT, or 1 μM ROT alone or in conjunction with 1 μM MLi2. **F.** ROS quantification using DHE signal intensity in WT 293-T cells. While co-treatment with MLi2 significantly reduced ROS from exposure to 500 nm ROT (p=0.0048), it could not attenuate ROS resulting from 1 μM ROT treatment (p=0.2740).One-way ANOVA with Tukey post-hoc for multiple comparisons, N = 3 – 4 biological independent replicates, 150 - 200 cells per technical replicate, *p<0.05, **p<0.01, ***p<0.001, ****p<0.0001.

### 3.3. LRRK2 inhibition attenuates toxicant-induced deficits in mitophagy

Mitophagy, a selective form of autophagy targeting damaged mitochondria and partially negatively regulated by LRRK2^18,32,37,38,46^ was assessed via quantification of mitophagy puncta in WT 293-T, G2019S, and LRRK2 KO cells treated with DMSO, ROT (500 nM) or TCE (500 μM), with or without 1 µM MLi2. The selection of these two toxicants was based on existing data showing that systemic administration of ROT or TCE induces the selective degeneration of dopaminergic neurons in models of parkinsonism caused by mitochondrial dysfunction^20,47–49^. Mitophagy was defined as the intersection of LC3b (lipidated autophagosomal membrane marker, green) with TOM20 (outer mitochondrial membrane marker, red; **Fig. 3A**), with a 3D volumetric reconstruction of puncta (**Supplemental Fig. 1A**). While some TOM20 changes were observed between cell groups (**Supplemental Fig. 1B**), the significant differences observed with just mitophagy puncta alone remained when controlling for mitochondrial number by normalization to TOM20 mean intensity (**Supplemental Fig. 1C**). Mitophagy analysis revealed that exposure to ROT and TCE significantly reduced the number of mitophagy puncta in WT 293-T and G2019S cells (**Fig. 3B-C**; WT ROT: p=0.0234, WT TCE: p=0.0008; G2019S ROT: p=0.0766, G2019S TCE: p=0.0003, two-way ANOVA). Co-treatment with MLi2 rescued the deficit in mitophagy puncta in both WT (WT 293-T ROT: p=0.0037, TCE: p<0.0001, one-way ANOVA) and G2019S mutant 293-T cells (G2019S ROT: p<0.0001, TCE: p<0.0001, one-way ANOVA), indicating that LRRK2 inhibition mitigates toxicant-induced mitophagy impairment (**Fig. 3C -3D**). Additionally, LRRK2 KO cells exhibited protection against the mitophagy deficits observed in both WT and G2019S cells, suggesting that LRRK2 plays a central role in toxicant-induced mitophagy disruption (**Fig. 3E**).

**Figure 3.**
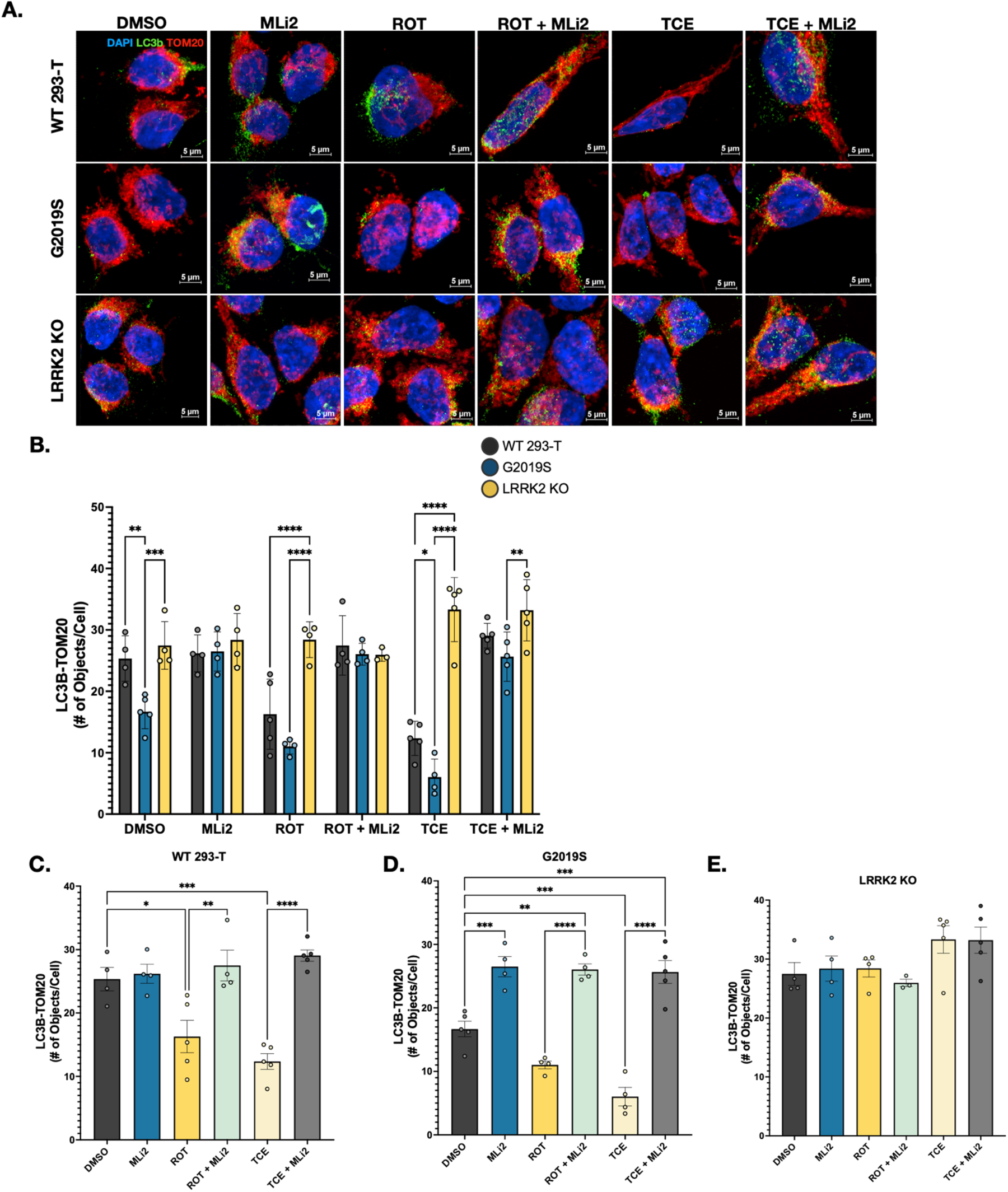
LRRK2 inhibition protects against toxicant-induced mitophagy deficits. **A.** Representative images of WT 293-T, G2019S, or LRRK2 KO treated with 0.10% DMSO, 1 μM MLi2, 500 nM ROT, and 500 μM TCE alone or in combination with 1 μM MLi2. Mitophagy was measured via a quantification of the number of intersecting objects between LC3b (autophagosomal membrane marker, green) and TOM20 (outer mitochondrial membrane marker, red). **B.** Quantification of number of LC3b/TOM20 intersecting objects per cell in each genotype. Two-way ANOVA revealed a significant interaction between genotype and treatment (F(10,60) = 9.612, p<0.0001), suggesting that LRRK2 plays a role in mitophagy deficits associated with toxicant exposure. Post-hoc Dunn’s multiple comparisons revealed genotype differences in DMSO, ROT, TCE, and TCE + MLi2 treated groups. **C-E.** Statistical comparisons between treatments in each individual genotype. Two-way ANOVA with Dunn’s multiple comparisons. MLi2 co-treatment reduced toxicant-induced mitophagy deficits for ROT (WT 293-T p=0.0037, G2019S p < 0.0001) and TCE (WT 293-T and G2019S p < 0.0001). LRRK2 KO cells were protected from mitophagy deficits and LRRK2 inhibition with MLi2 has no off-target effects. One-way ANOVAs with post-hoc Tukey’s multiple-comparisons. WT 293-T F(5,21) = 14.75, p <0.0001, G2019S F(5,20) = 38.63, p<0.0001, LRRK2 KO F(5,19) = 2.266, p = 0.0893. N = 3 – 4 biological replicates, n = 150 – 200 cells per technical replicate, *p<0.05, **p<0.01, ***p<0.001, ****p<0.0001.

### 3.4. LRRK2 kinase inhibition protects against TCE-induced neurodegeneration *in vivo*

We previously showed that TCE caused elevated LRRK2 kinase activity in the striatum and substantia nigra of rats after 3 weeks of exposure, prior to dopaminergic neurodegeneration. To identify if LRRK2 kinase inhibition was protective against TCE exposure in this *in vivo* model, adult (12-month-old) female Lewis rats were treated with 200 mg/kg TCE or vehicle (olive oil, oral gavage) with 10 mg/kg MLi2 or control (1% methyl cellulose) beginning after 3 weeks TCE exposure until 6 weeks (**Fig. 4A**). Stereological assessment of rat brain tissue confirmed that TCE and VEH treated rats had no significant loss of dopaminergic neurons in the SN at 3 weeks (**Fig. 4B-C**). After 6 weeks of TCE exposure however, rats displayed ∼45% loss of dopaminergic neurons in the substantia nigra compared to VEH, and that post-lesion treatment with MLi2 significantly protected against cell loss (one-way ANOVA, p-value<0.0001, Tukey’s post-hoc multiple comparisons; **Fig 4D-F**). Nissl staining of midbrain tissue sections confirmed the loss of Nissl-positive cells in the SN of TCE-treated animals, while post-lesion treatment with MLi2 prevented cell loss (p value<0.0001, one-way ANOVA; **Fig 4E-G**).

**Figure 4.**
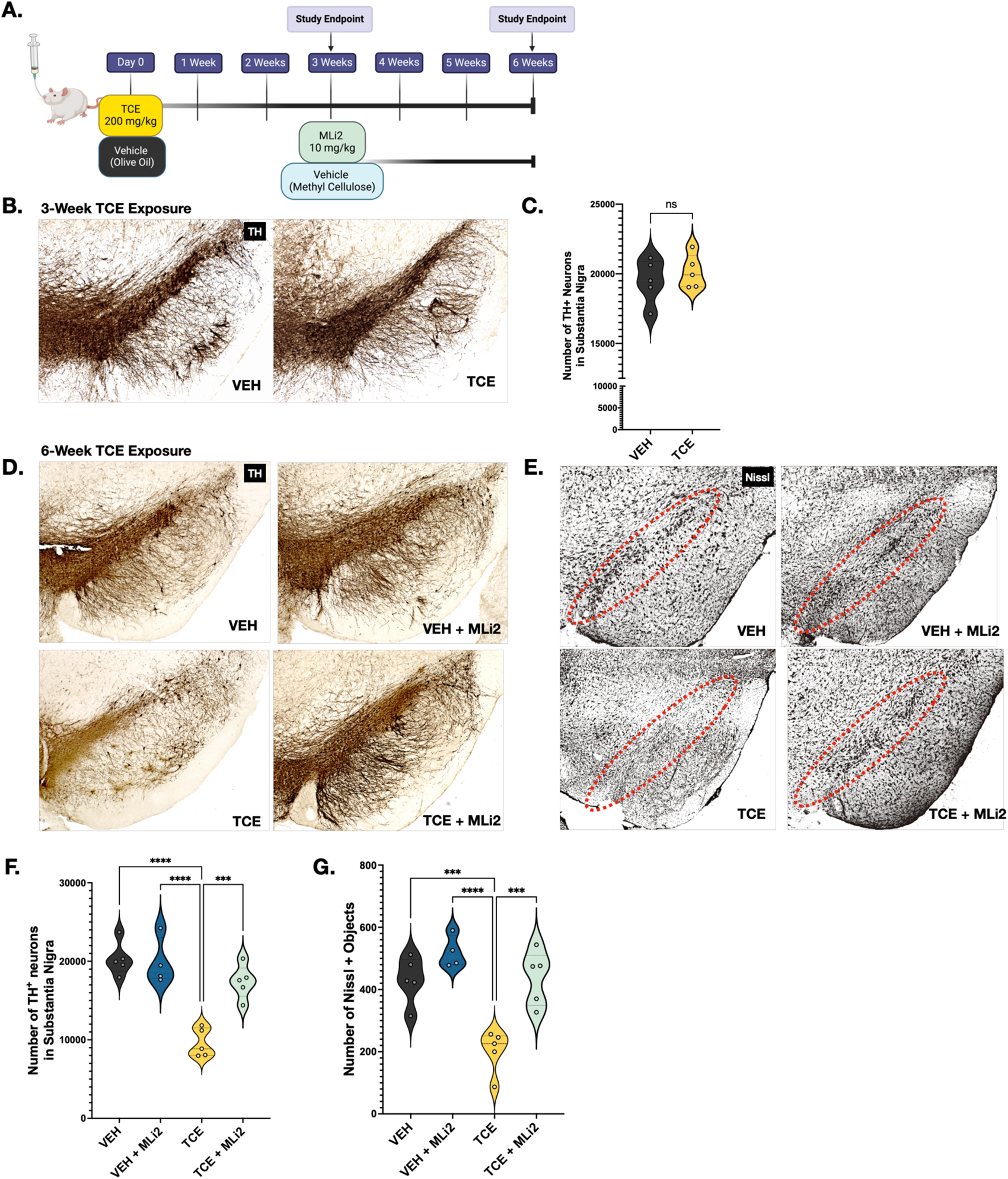
LRRK2 kinase inhibition attenuates TCE-induced neurodegeneration. **A.** Aged female Lewis rats were treated with 200 mg/kg of TCE for 6 weeks. Beginning at 3 weeks post-exposure, a cohort of rats was treated with 10 mg/kg MLi2 or vehicle**. B-C.** TH-positive cells from 3-week VEH and TCE exposed rats within the SN, N=5 animals per group. Unpaired Mann-Whitney U Test (p=0.5952) indicates no differences between TCE and VEH animals at 3 weeks. **D.** Representative images of Nissl-stained VEH and TCE-exposed mice at 6 weeks with SN highlighted. **E.** Representative images of TH-positive cells within the SN of VEH or TCE-treated rats for 6 weeks. **F.** Stereological counts of TH+ cells in each group. MLi2 post-lesion administration significantly rescued neuronal loss in TCE-treated animals, p=0.0003 (F=23.47, p<0.0001). **G.** Analysis of Nissl+ objects in the SN. MLi2 post-lesion administration significantly rescued cell loss in the SN in TCE-treated animals, p=0.0007 (F=16.40, p<0.0001). ANOVA tables, descriptive statistics, and post-hoc comparisons are provided in Supplemental Table 1. N = 4-5 animals per group. Violin plots represent the data distribution with its probability density, with dashed lines representing mean. Significance values as indicated: ***p<0.001, ****p<0.0001.

### 3.5. LRRK2 kinase inhibition attenuates oxidative stress in dopaminergic neurons of TCE-exposed rats

To investigate the contribution of LRRK2 to oxidative stress *in vivo*, oxidative stress in DA neurons in the SN was measured using markers for oxidative protein modification after 6-weeks of exposure. Significantly elevated levels of 3-nitrotyrosine (3-NT), a marker for peroxynitrite (ONOO^−^) protein adducts on tyrosine residues were observed in DA neurons of TCE-exposed rats (p=0.0002), which were significantly attenuated by MLi2 post-lesion treatment (p=0.0003; **Fig 5A, C**). DA neurons of the SN in TCE-treated animals also showed elevated 4-hydroxynonenal (4-HNE; p<0.0001), a marker for lipid peroxidation caused by oxygen radicals, which was significantly rescued via LRRK2 kinase inhibition (p < 0.0001; **Fig 5B, D**). Together, these data indicate surviving dopaminergic neurons in TCE-treated animals have elevated levels of oxidative damage, and LRRK2 kinase inhibition significantly decreased TCE-induced oxidative damage within these neurons, potentially as a mechanism of neuroprotection.

**Figure 5.**
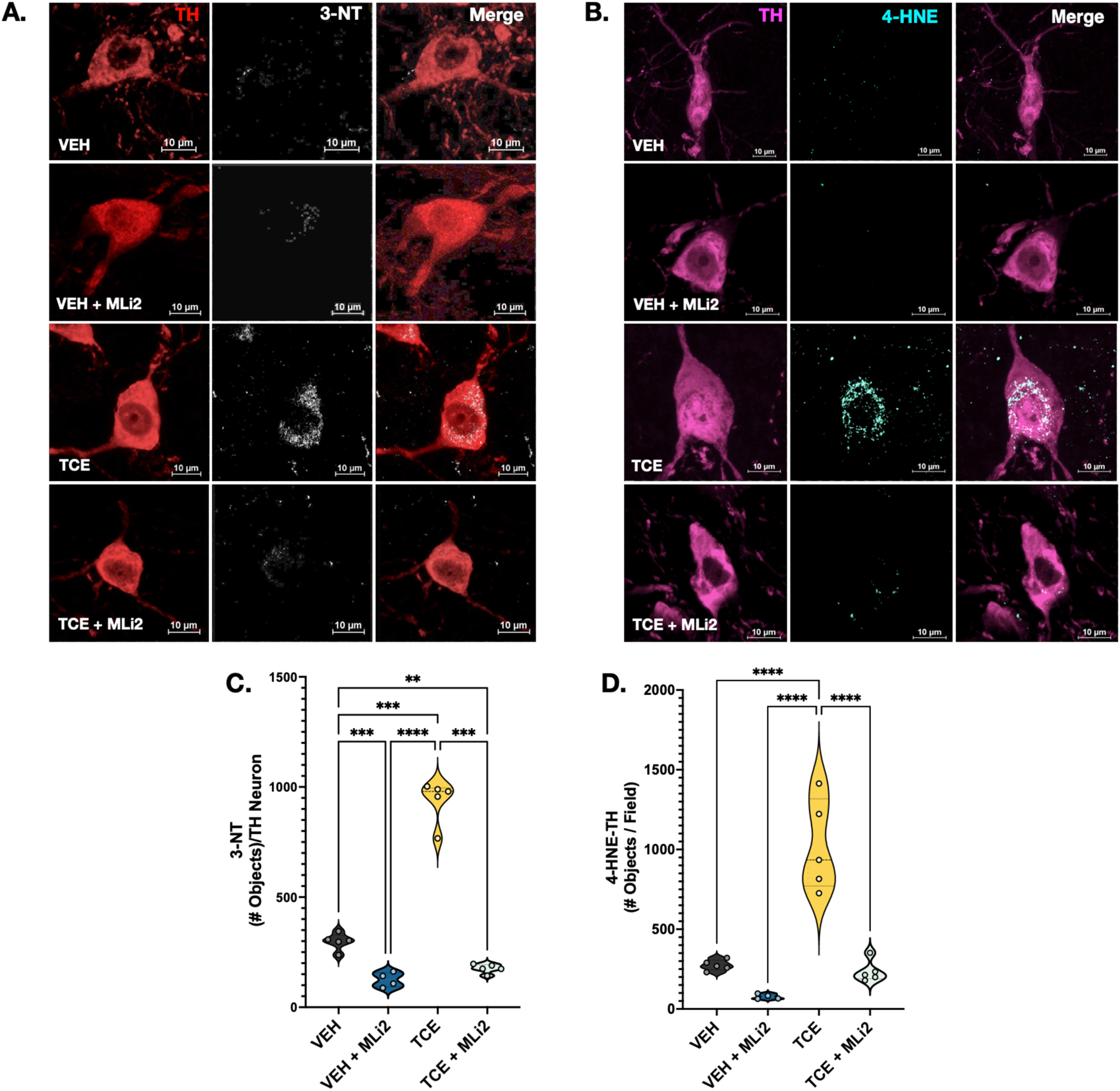
LRRK2 kinase inhibition reduced oxidative stress in dopaminergic neurons of TCE-exposed rats. **A-B**. Representative images of dopaminergic neurons (60x) within the SN of VEH of TCE-exposed animals for 6 weeks, with or without post-lesion treatment with 10 mg/kg MLi2. Oxidative damage was measured using 3-nitrotyrosine (3-NT, white) and 4-hydroxynonenal (4-HNE) cyan. **C.** Quantification of 3-NT per TH+ neuron. TCE animals treated with post-lesion administration of MLi2 had reduced oxidative stress compared to TCE animals with vehicle (methyl cellulose), p=0.0003, F(3.00, 6.734)=219.9, p<0.0001, Brown-Forsythe ANOVA with post-hoc Dunnett’s for multiple comparisons). **D.** Quantification of 4-HNE per TH+ SN field. MLi2 treatment significantly reduced oxidative stress in DA neurons of TCE animals at 6 weeks (p<0.0001) with significant differences in oxidative stress from the peroxidation of fatty acids (F(3,15)=36.04, p<0.0001, one-way ANOVA with Tukey post-hoc multiple comparisons). N=4-5 animals per group. ANOVA tables, descriptive statistics, and post-hoc comparisons are provided in Supplemental Table 1. Violin plots represent the data distribution with its probability density, with dashed lines representing means of groups. Significance values as indicated: ***(p<0.001), ****(p<0.0001).

### 3.6. LRRK2 kinase inhibition prevented mitochondrial damage in TCE-treated animals

To evaluate the role of LRRK2 in mitochondrial damage in DA neurons in the SN of TCE-treated rats, brain tissue from animals exposed for 6 weeks was stained for TH (red), pS65-Ub (green) a ubiquitin residue specific to damage-mediated pexophagy and mitophagy, and TOM20 (magenta); **Fig 6A-B**). The size-restricted intersection of pS65-Ub and TOM20 within TH+ neurons was analyzed per field, as visualized in **Fig 6B**, to provide a metric for damaged mitochondria in dopaminergic neurons. Statistical analyses revealed that TCE-treated animals had greater quantities of uncleared damaged mitochondria compared to control animals (p=0.0005), which were decreased with administration of MLi2 (p<0.0001; **Fig 6D**).

**Figure 6.**
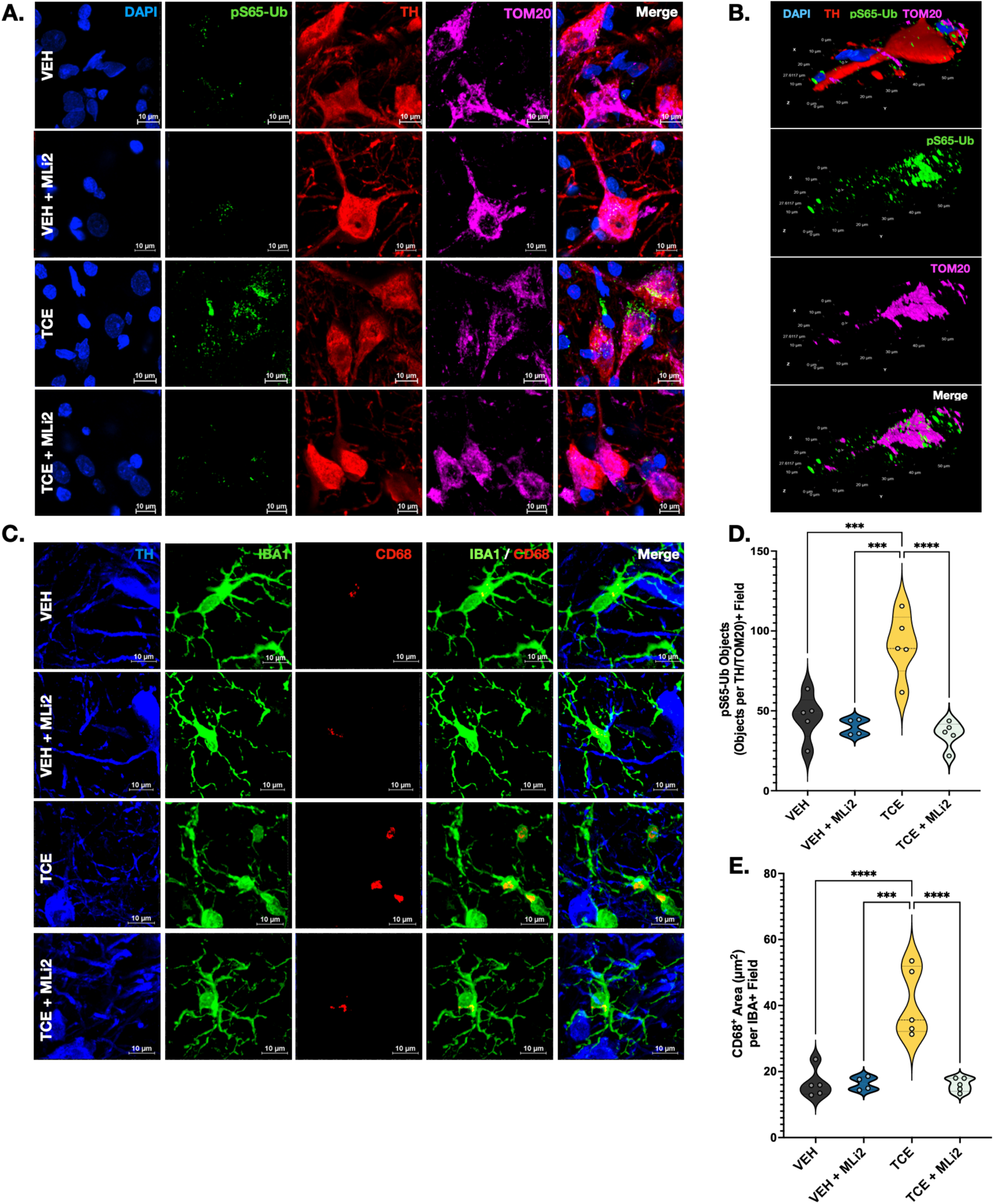
LRRK2 kinase inhibition reduced mitochondrial damage and neuroinflammation from TCE exposure. **A**. Representative images of ps65-Ub, TOM20, and TH in DA neurons of SNPc show a reduction of mitochondrial damage in TCE-treated animals with post-lesion LRRK2 kinase inhibition at 6 weeks. **B.** Volumetric re-composition of a DA neuron in SNPc of a TCE-treated animal demonstrating the intersection of TOM20 and pS65-Ub within the neuron. Scale bars are depicted on the side of each plane. **C.** Representative images of CD68, IBA1, and TH in SN. **D.** Quantification of ps65-Ub intersecting objects per TH/TOM20+ field in SNPc demonstrated differences in mitochondrial damage with MLi2 treatment at 6 weeks (F(3, 15)=17.89, p<0.0001, one-way ANOVA with post-hoc Tukey’s for multiple comparisons). TCE animals treated with MLi2 had reduced mitochondrial damage compared to TCE animals treated with vehicle (methyl cellulose), p<0.0001. **E.** Quantification of CD68+ objects per IBA1 field demonstrated significant differences in microglial activation with MLi2 treatment (F(3,15)=20.72, p<0.0001, one-way ANOVA with Tukey post-hoc multiple comparisons). MLi2 treatment significantly reduced microglial activation in SNPc of TCE-exposed animals compared to TCE animals treated with methyl cellulose at 6 weeks, p<0.0001. N=4-5 animals per group. ANOVA tables, descriptive statistics, and post-hoc comparisons are provided in the Supplemental Table 1. Violin plots represent the data distribution with its probability density, with dashed lines representing means of groups. Significance values as indicated: ***(p<0.001), ****(p<0.0001).

### 3.7. TCE-induced microglial activation is reduced by LRRK2 kinase inhibition

We previously showed activated microglia within the SN of TCE-exposed rats, which could be related to mitochondrial dysfunction and oxidative stress caused by TCE, and possibly modulated by LRRK2^22^. To assess this, SN sections from 6-week animals were stained for TH, IBA1, and CD68, a lysosomal protein indicative of phagocytic activity (**Fig 6C**). We observed a significant elevation of CD68 in microglia of the SN of TCE-treated animals after 6 weeks of exposure (p<0.0001), which was attenuated by treatment with MLi2 (p<0.0001, **Fig 6E**).

## Discussion

Both genes and environmental factors contribute to PD risk, and most cases likely stem from the interaction between them (GxE). Genetic and environmental risk factors converge on several cellular pathways that influence neurodegeneration in PD, such as mitochondrial dysfunction, endolysosomal deficits, and inflammation^17,26,32,50^. Here, we identify that WT LRRK2 plays a key role in toxicant-induced dopaminergic pathology, and its inhibition could serve as a protective mechanism against neurotoxicity caused by environmental risk factors for PD. Furthermore, we found that mutations in LRRK2 may augment the effects of environmental risk factors, such as mitochondrial toxicants, which could explain some of the heterogeneity in PD phenoconversion in individuals with pathogenic LRRK2 variants.

We and others have previously shown that environmental toxicants, such as rotenone and TCE, elevate WT LRRK2 kinase activity in the brain to a similar degree as the most well characterized pathogenic LRRK2 mutation, G2019S^22,25,51,52^. Furthermore, growing data suggests that LRRK2 moderates PD-related pathology caused by exposure to toxicants, such as microglial inflammation induced by the heavy metal manganese^35,53,54^. Conversely, both pharmacologic and genetic LRRK2 inhibition are protective against neuropathology caused by environmental risk factors for PD, including a reduction in brain inflammation caused by paraquat^55^, improved survival of manganese exposed macrophages^53,54^, and neuroprotection of dopaminergic neurons in a rotenone model of PD^25^. Here, we similarly demonstrated that pharmacologic inhibition of LRRK2 by the small molecule MLi2 was protective in a model of TCE-induced neurodegeneration we previously characterized showing elevated nigrostriatal LRRK2 kinase activity before the loss of dopaminergic neurons^23^.

Toxicants associated with PD risk vary in structure, mechanism of action, and potency. For example, the organic pesticide rotenone is a large (394.41 g/mol) crystalline isoflavone that acts as a prototypical complex I inhibitor^56,57^. In contrast, TCE is a small (131.4 g/mol) chlorinated solvent that is reported to inhibit mitochondria at high doses^47,58^, possibly through its metabolites such as S-(1,2-dichlorovinyl)-L-cysteine (DCVC)^15^. Similarly, PERC (165.83 g/mol) appears to decrease electron flow at the inner mitochondrial membrane^19^, and paraquat (257.16 g/mol) is characterized as an inducer of redox cycling resulting in toxic ROS production and permeability of the inner mitochondrial membrane^6,59,60^. Thus, despite differences in precise mechanism, each toxicant associated with PD risk causes direct or indirect mitochondrial dysfunction^13,19,25,47,49,60^, which likely acts as a common mechanism associated with parkinsonian neurotoxicity. This was consistent with our findings that each toxicant, when used at subtoxic doses in cell treatments, caused a similar degree of ROS generation over a spectrum of time; early/immediate for rotenone and paraquat and delayed for TCE and PERC, consistent with the data suggesting a metabolite of the solvents ultimately causes mitochondrial toxicity.

As a key convergence point, we found that regardless of the toxicant, WT LRRK2 played a role in oxidative stress as LRRK2 KO cells or cells treated with MLi2 were markedly protected from toxicant-induced ROS, but this effect was exacerbated in cells with the G2019S mutation. Given the role of LRRK2 in endolysosomal trafficking and its link to mitophagy^37,38,61–63^, we suspected that toxicant-damaged mitochondria may have impaired clearance in cells with aberrantly activated LRRK2 and that LRRK2 inhibition promotes the removal of damaged mitochondria via mitophagy^18,37,38^. *In vitro*, we observed this with the strong loss of colocalization of LC3b with TOM20 in cells exposed to rotenone or TCE, which was ameliorated by MLi2 co-treatment and protected in LRRK2 KO cells. In parallel, elevated pS65-Ub, a marker of mitochondrial damage, was elevated in dopaminergic neurons of TCE-exposed animals, which was significantly reduced by LRRK2 inhibition. Although dynamic measures of mitophagy cannot be measured in fixed brain tissue, as the *in vivo* TCE study was performed using MLi2 “post-lesion,” beginning 3 weeks after the start of TCE exposure when LRRK2 kinase activity is elevated^23^, we predict that mitochondrial damage caused by TCE was at least partially ameliorated. This is further supported by the significant reduction in oxidative damage measured with 3-NT and 4-HNE after 6 weeks of TCE exposure in MLi2-treated mice, particularly given the perinuclear appearance of these oxidative residues, which coincide with the expression of pS65-Ub and TOM20 at the mitochondria.

The mechanisms by which LRRK2 inhibition protects against neurodegeneration caused by toxicant exposure has been a key question as these model systems partially recapitulate the most common form of idiopathic PD, without genetic mutations^17,26,32,64^. As LRRK2 inhibitors continue their development through clinical trials, understanding the mechanisms that underlie the pathological contribution of WT, endogenously expressed LRRK2 may provide evidence for the broader use of LRRK2 inhibition as a therapeutic strategy against PD. The data from this study suggest that while the LRRK2 G2019S mutation did exacerbate toxicant-driven ROS and reduced mitophagy, inhibition of *wildtype* LRRK2 was sufficient to protect against all measures of toxicity caused by each mitochondrial toxicant tested *in vitro* and was ultimately protective against dopaminergic neurodegeneration *in vivo*. Conversely, though the mutation is relatively rare at the population level, exposure to environmental mitochondrial toxicants could be the trigger necessary to induce neurodegeneration in the incompletely penetrant G2019S polymorphism^34^.

Despite the near complete protection against neurotoxicity in our experimental systems here, LRRK2-mediated effects cannot explain every pathological endpoint associated with environmental toxicants implicated in PD risk^65,66^. For example, we found that a threshold exists for LRRK2 protection against mitochondrial toxicity. Using rotenone as a proof-of-principle complex I inhibitor, LRRK2 KO cells were protected against ROS caused by low-dose (500 nM) rotenone, but this protection did not continue at doses higher than 1 µM. Likewise, WT cells were no longer protected from 1µM rotenone by MLi2. As 1 µM rotenone is likely approaching near complete complex I inhibition^67^, we suspect that overt toxicity overwhelms the protective effect of LRRK2-mediated effects such as improved mitochondrial clearance. However, as most exposures associated with PD likely happen at low doses over chronic periods^1,4,7,8,64,68,69^, LRRK2 could be involved in crosstalk with pathways triggered in parallel by environmental risk factors^26^. For example, mitochondrial damage from toxicants can lead to pro-inflammatory cascades via recruitment of cytosolic sensors such as the cyclic GMP-AMP synthase activator and stimulator of interferon genes (cGAS-STING)^70–74^. As LRRK2 is highly expressed in immune cells, such as macrophages^75–77^, and is implicated in several critical processes in inflammation^35,54,55^ such as cytokine release, autophagy, and phagocytosis^32,78,79^, LRRK2 may provide a link between toxicant-induced mitochondrial dysfunction and inflammation in PD. In support of this, we showed here that LRRK2 inhibition reduced microglial activation in rat brain tissue, which coincided with a reduction in mitochondrial damage caused by TCE exposure. Similarly, in Rocha et al., 2019, we and our collaborators showed that LRRK2 inhibition rescued inflammation and endolysosomal dysfunction caused by rotenone^63^. Thus, aberrantly activated LRRK2 associated with environmental toxicants may influence direct oxidative damage through mitochondrial dysfunction and perpetuate chronic effects such as inflammatory signaling cascades and protein accumulation that continue after exposure has ceased.

While the data here represent several key mechanisms that explain how LRRK2 may partially contribute to toxicant-induced neurodegeneration in PD, there are also limitations that need to be considered. First, the expression levels of endogenous WT LRRK2 necessitated using a HEK cell line that expresses LRRK2 at measurable levels^26,52^. While different from neurons, HEK cells are extensively characterized in the LRRK2 literature^25,26,44,80,81^, have been used as a model system to study the effects of LRRK2 mutations and display similar energetic demands of neurons that rely heavily on mitochondrial function^82–84^. We also strategically chose to evaluate toxicants linked to PD risk with structural dissimilarities but associated with mitochondrial dysfunction^13,19,47,56–60,85^. However, we could not feasibly compare each in an *in vivo* model. To this end, we selected a model of TCE exposure where we previously characterized elevated LRRK2 kinase activity preceding neurodegeneration in the nigrostriatal tract over time^23^. We have recently worked to develop a more environmentally relevant TCE exposure system involving passive inhalation^43^; however, we do not currently have the same level of LRRK2 kinase activation data as this model extends over a chronic period. In our ongoing studies, we will continue to evaluate the role of LRRK2 in TCE-induced neurodegeneration using an environmentally relevant dose and route of exposure to further uncover the mechanisms linking PD-related toxicants and LRRK2.

## Acknowledgements

This work was funded by the National Institute of Environmental Health Science (R00ES029986, BRD), and the UAB Training Program in Neurodegeneration (T32NS095775, NMI).

---

**Supplemental Figure 1.**
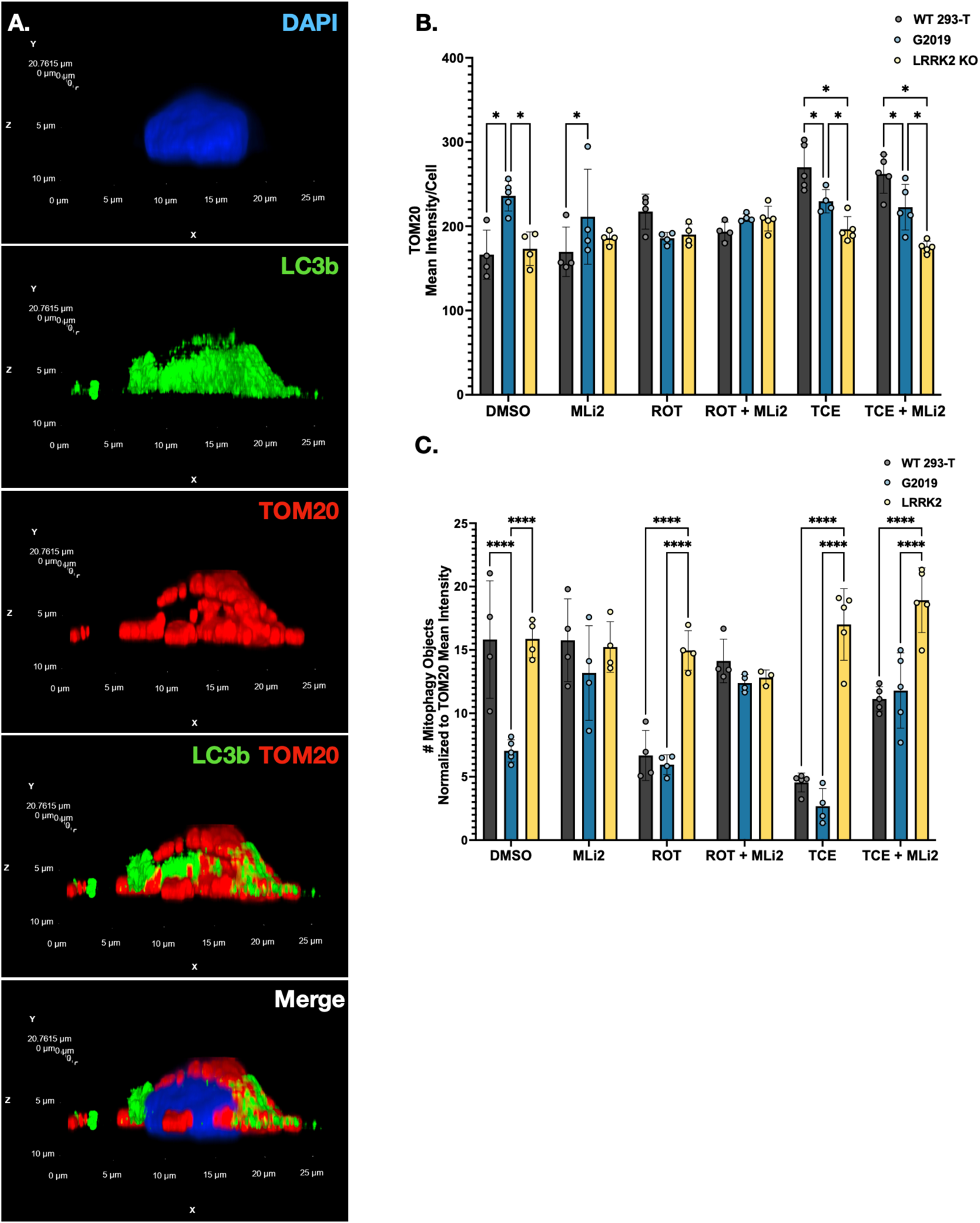
Normalization to TOM20 demonstrated no alteration in mitophagy findings. **A**. Volumetric re-composition of a LRRK2 KO 293-T HEK cell treated with 500 nM ROT and stained with TOM20, LC3b, and DAPI. Scale bars are depicted on the side of each plane. **B.** Quantification of TOM20 mean intensity/cell for all mitophagy data analyzed demonstrated effects of genotype and treatment in TOM20 mean intensity per cell (F(10, 61) = 7.636, p<0.0001, two-way ANOVA). **C.** Quantification of mitophagy puncta (LC3b/TOM20 intersecting objects) normalized to TOM20 mean intensity demonstrated significant effects of both treatment and genotype (F(10, 59) = 9.764, p<0.0001, two-way ANOVA). ANOVA tables, descriptive statistics, and post-hoc comparisons are provided in Supplemental Table 1. N = 3 - 4 independent biological replicates, 120 - 160 cells per technical replicate. Bar graphs represent mean values for each group, with error bars. Significance values as indicated: * (p<0.05), ****(p<0.0001).

